# Additive genetic variance for lifetime fitness and the capacity for adaptation in an annual plant

**DOI:** 10.1101/601682

**Authors:** Mason W. Kulbaba, Seema N. Sheth, Rachel E. Pain, Vince M. Eckhart, Ruth G. Shaw

## Abstract

The immediate capacity for adaptation under current environmental conditions is directly proportional to the additive genetic variance for fitness, V_A_(W). Mean absolute fitness, 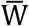, is predicted to change at the rate 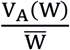, according to Fisher’s Fundamental Theorem of Natural Selection. Despite ample research evaluating degree of local adaptation, direct assessment of V_A_(W) and the capacity for ongoing adaptation is exceedingly rare. We estimated V_A_(W) and 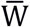 in three pedigreed populations of annual *Chamaecrista fasciculata,* over three years in the wild. Contrasting with common expectations, we found significant V_A_(W) in all populations and years, predicting increased mean fitness in subsequent generations (0.83 to 6.12 seeds per individual). Further, we detected two cases predicting “evolutionary rescue”, where selection on standing V_A_(W) was expected to increase fitness of declining populations (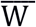 < 1.0) to levels consistent with population sustainability and growth. Within populations, interannual differences in genetic expression of fitness were striking. Significant genotype-by-year interactions reflected modest correlations between breeding values across years (all *r* < 0.490), indicating temporally variable selection at the genotypic level; that could contribute to maintaining V_A_(W). By directly estimating V_A_(W) and total lifetime 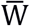, our study presents an experimental approach for studies of adaptive capacity in the wild.

## Introduction

A population’s mean absolute fitness, 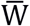, in terms of the per capita contribution of offspring, corresponds to its growth rate and is thus a measure of its degree of adaptation (Fisher 1930, pp. 25, 37; Roughgarden 1996, Ch. 4). Fisher (1930) showed that the immediate capacity for further adaptation is proportional to the magnitude of a population’s additive genetic variance for absolute fitness, evaluated as an individuals’ lifetime contribution of offspring to the population (Fisher 1930; Price 1970, 1972; Ewens 2004). Moreover, this Fundamental Theorem of Natural Selection (FTNS, Fisher 1930) quantitatively predicts the rate of adaptation under prevailing environmental conditions as the ratio of V_A_(W) to current mean population fitness, 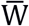. This ratio predicts the genetically based change, from one generation to the next, in the per capita rate of population growth. Such predictions are vital to understanding the adaptive process, by first determining the possibility for evolutionary change in mean fitness, and second, the expected magnitude of fitness increase in the wild. Thus, using FTNS to predict future adaptation under ongoing natural selection requires direct, quantitative estimates of standing additive genetic variance for fitness, V_A_(W). Studies that take such a direct and predictive approach are exceedingly rare (but see Etterson 2004a,b; Winn 2004; Sheth et al. 2018).

Studies of ongoing selection have generally focused on traits of particular interest. Lande and Arnold’s (1983) proposal of multiple regression of fitness on traits powerfully stimulated research of this kind. As they noted, this approach reflects only the selection on those traits under consideration, through their associations with the measure of fitness. Few such studies report the extent to which the included traits account for the variation in fitness (but see Conner 1988; Janzen 1993; Conner et al. 1996a,b). Natural selection likely bears on many more traits, however, and this implies that the included traits typically account for a modest proportion of the variation in fitness (see Shaw 2019). Whereas the regression of fitness on a set of traits can yield insight into evolutionary change in particular traits under selection, they are ill-suited for predicting the change in mean absolute fitness, i.e. the degree of overall adaptation.

Retrospective studies of genetic differentiation and adaptation to changes in local conditions have amply demonstrated the efficacy of selection in the past. These cases are illuminating even when the aspect(s) of the environment that impose(d) differential selection are unknown, as is often the case, given the high dimensionality of environment. When the timing of an abrupt environmental change is known, the rate that adaptation proceeded can also be determined, or at least bounded. Cases of rapid adaptation, within a few to tens of generations, are now well known for a variety of taxa and include adaptation that followed changes in edaphic conditions (Antonovics and Bradshaw 1970; Al-Hiyaly et al. 1993), introduction of herbicides (Vigueira et al. 2013; Baucom 2019), predators (Fisk et al. 2007), and changing climate (Franks et al. 2007; Geerts et al. 2015). These examples of rapid adaptation to local conditions reveal capacity for adaptation to environmental differences from standing genetic variation (Barrett and Schluter 2008). However, such studies are limited to characterizing the current degree of adaptation in response to past selection and do not evaluate the potential for ongoing adaptation.

Quantitative prediction of the rate of adaptation from FTNS is vital to understanding ongoing adaptation in wild populations, especially in assessing whether standing levels of V_A_(W) suffice for contemporary natural selection in declining populations (i.e. 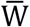 < 1) to halt further decline (Bradshaw 1991). Such “evolutionary rescue” (Gomulkiewicz and Holt 1995) that increases fitness to 1 (i.e. individuals demographically replacing themselves) or greater depends on current standing V_A_(W). However, a sufficient V_A_(W) for such fitness increase is unknown, particularly in wild populations subject to substantial spatial and temporal environmental variation (Carlson et al. 2014). In the current context of drastic ongoing environmental change, estimates of the magnitude of a populations’ V_A_(W) are critical for predicting the genetically-based change in 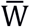 in response to the natural selection impinging on them. Given the elevated threat to population persistence due to rapid change in climate and other aspects of the environment (Shaw and Etterson 2012; Neumann et al. 2017), it is now imperative to clarify the efficacy of selection in maintaining and enhancing population mean fitness *in situ.*

Despite their importance, very few direct estimates of standing V_A_(W) exist (but see Etterson 2004a,b; Winn 2004; Sheth et al. 2018). Several factors may explain this paucity of empirical research. First, population mean fitness is often thought to be at or near optimum due to incessant selection, which would tend to fix or eliminate alleles having directional effects on fitness. Accordingly, V_A_(W), and therefore the capacity for adaptation, is conjectured to be reduced to low or negligible levels (e.g., Mazer 1987; Barton and Keightley 2002). However, even with consistent and strong selection at individual loci, fixation requires hundreds of generations (e.g., Fig. 6.3 in Hartl and Clark 1997; Messer et al. 2016). The continuing response to selection in the Illinois corn selection experiment (Moose et al. 2004), even after 100 generations, demonstrates the persistence of variation supporting a response to artificial selection on biochemical constituents of kernels. However, the resilience of variation under selection in nature remains an open empirical question. Second, numerous studies have found that estimates of narrow-sense heritability 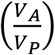 for individual fitness components tend to be lower than for morphological traits (Roff and Mousseau 1987; McFarlane et al. 2014). Because environmental variation inherent to fitness components is compounded with environmental variance of underlying morphological traits, high environmental variance, rather than low additive genetic variance, may explain low heritability of fitness components (Price and Schluter 1991). Nevertheless, the empirical generalization about lower heritability of individual components of fitness contributed to the expectation that additive genetic variance for lifetime fitness is likely to be modest. This expectation may, however, be inaccurate because it does not account for the dependence of sequential fitness components through the life-span. Third, when pedigrees are not experimentally derived and may be uncertain (e.g. Foerster et al. 2007), logistical challenges encumber efforts to both reconstruct relatedness of individuals and determine total lifetime fitness. Uncertainty in the assignment of relatedness casts doubt on the estimates of quantitative genetic parameters (Thomas et al. 2000). Finally, confounding between genetic and environmental influences, likely in observational studies, can also bias V_A_(W) estimates (Pemberton 2010), and therefore inferences drawn from them.

Whereas direct estimates of V_A_(W) in the wild are rare, theory bearing on whether substantial genetic variation could be maintained is ambiguous. In particular, early theory addressing the role of environmental variation, particularly temporal variability, indicated that restrictive conditions are required to maintain genetic variation within populations (Felsenstein 1976; Hedrick 1986; Gillespie and Turelli 1989). More recently, models focusing on V_A_(W) under environmentally varying selection have demonstrated maintenance of substantial V_A_(W) (Zhang 2012; Shaw and Shaw 2014; see also Yi and Dean 2013).

Direct estimation of V_A_(W) is not a trivial endeavor, requiring sound experimental design and statistical inference. Experimental approaches that facilitate estimates of complete lifetime fitness expressions from fully pedigreed individuals, and avoid confounding environmental and genetic effects are essential (Shaw 2019). Further, statistical tools are needed that can validly model lifetime fitness, which conforms to no conventional statistical distribution (Shaw et al. 2008). Aster models account for the dependent nature of sequential life-history fitness expressions to estimate total mean lifetime fitness (Geyer et al. 2007), and random parental effects (Geyer et al. 2013) to estimate the additive genetic effects on lifetime fitness.

In this study, we estimate the capacity for adaptation under contemporary natural selection in wild populations of the annual legume *Chamaecrista fasciculata* growing in their native locations. Using Fisher’s Fundamental Theorem (Fisher 1930) and aster modelling to analyze records of lifetime fitness (Geyer et al. 2007, 2013), we directly estimate V_A_(W) and 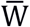 to quantitatively predict the rate of adaptation. To assess the generality of our findings, we conducted our experiment in three populations in three successive years. Our experimental design allowed us to characterize population-specific expression of V_A_(W) and to predict the rate of future adaptation, while also evaluating temporal variation in the expression of V_A_(W), and therefore variation in the capacity for adaptation, in three populations over three years.

## Materials and Methods

### Study system

We estimated V_A_(W) and 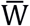 as the basis for predicting the generational change in mean fitness in three populations (Fig. 1) of the annual legume *Chamaecrista fasciculata*, each growing in its home site. The range of *C*. *fasciculata* spans from the prairies of the Northern Great Plains in Minnesota to Central Mexico and to the eastern seaboard of North America (Irwin and Barneby 1982). *C. fasciculata* produces hermaphroditic, enantiostylous flowers that are buzz-pollinated (Lee and Bazzaz 1982) by bumblebees *(Bombus* spp). Its experimental tractability has made it the focus of previous studies of gene flow (Fenster 1991a,b), quantitative genetics (Kelly 1993; Etterson 2004b), selection on traits (Etterson 2004a), local adaptation (Galloway and Fenster 2000), adaptation to climate change (Etterson and Shaw 2001; Etterson 2004b), evolution of range limits (Stanton-Geddes et al. 2012b, 2013), and genotype-by-environment interactions (Sheth et al. 2018).

**Figure 1.**
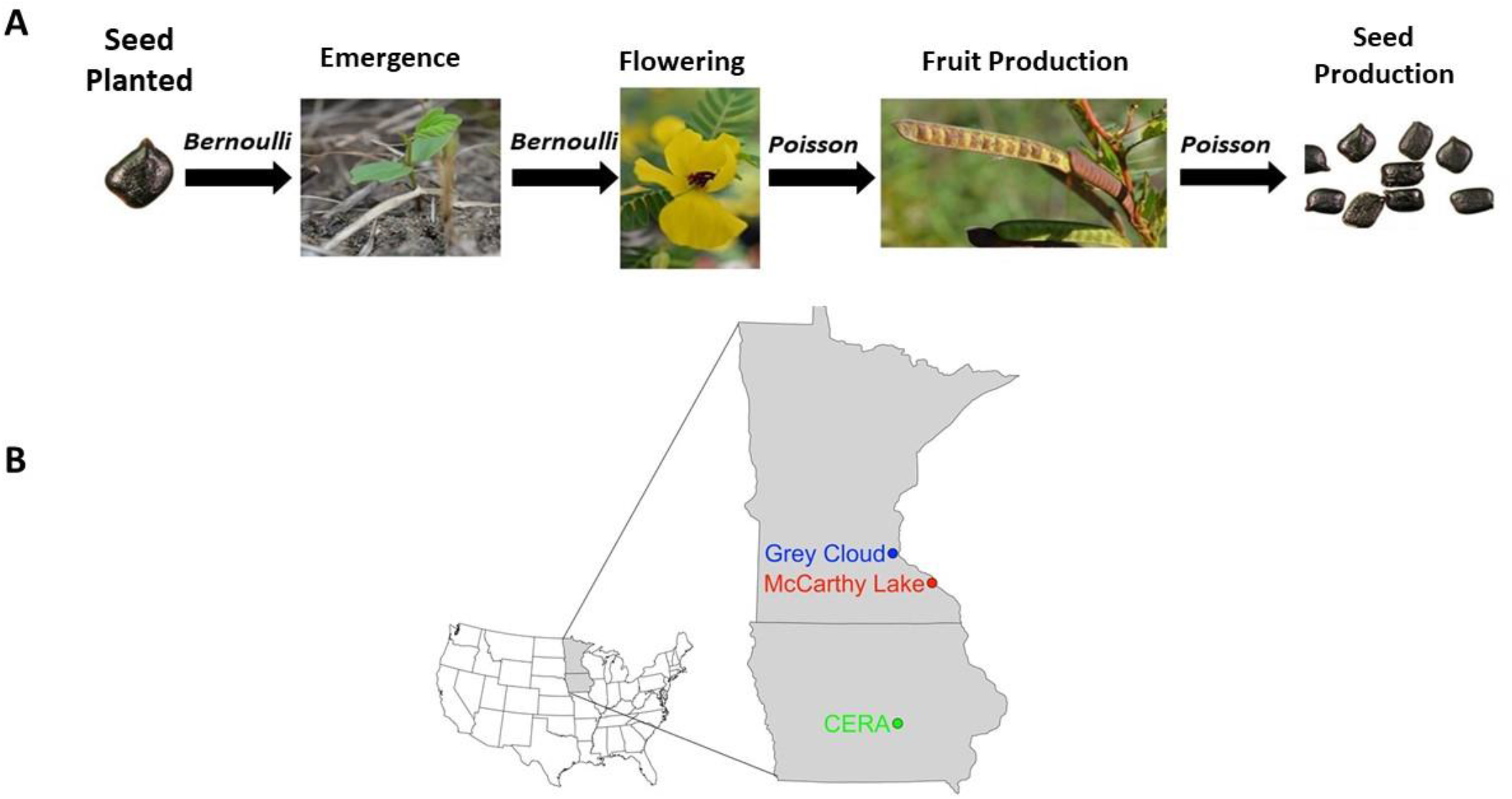
(A) Graphical model used to estimate lifetime fitness. Each node represents a fitness component and therefore response variable, and arrows represent conditional distributions. Probability of germination and flowering (0 or 1; Bernoulli distribution), and total number of fruits and seeds produced (Poisson distribution). Additional node to account for subsampling of total fruit production (not shown) also followed a Poisson distribution. (B) Map of three experimental populations. Seed photos from Anna Peschel and remaining photos by MWK.

### Genetic sources and pedigree design

In the fall of 2012, 2-3 fruits from 200 maternal plants no closer together than 10 m were collected from a population of *C*. *fasciculata* occupying remnant prairie at Kellogg-Weaver Dunes in Minnesota (44°15’43” N, 91°55’15” W; hereafter “McCarthy Lake”). Similarly the following fall, fruits from 200 maternal plants were collected from populations of *C*. *fasciculata* at Grey Cloud Dunes Scientific and Natural Area in Minnesota (44°46’32”N, 93°01’38”W; hereafter “Grey Cloud”), and 100 fruits from plants at the Conard Environmental Research Area in Iowa (41°40’44.2”N 92°51’24.9”W; hereafter “CERA”).

To obtain pedigreed populations of seeds from each of these populations, six seeds from each maternal plant were surface sterilized using an 8.9% bleach solution followed by a 70% ethanol rinse. We used 100 grit sandpaper to scarify the seeds, imbibed them in sterile water for three days, and then planted them in small peat pots. A total of 167 Grey Cloud, 196 McCarthy Lake, and 84 CERA individuals were used to generate pedigreed plants. One seedling per individual was used to represent each distinct field-collected family and was transplanted into a large tree pot with 1 teaspoon of Osmocote^®^ slow release fertilizer (14:14:14) after producing five true leaves. Plants were grown in the University of Minnesota Plant Growth Facilities greenhouse under 16:8-h photoperiod. Hand crosses were made according to a nested paternal half-sibling design (North Carolina Design I; Comstock and Robinson 1948), with independent sets of three dams randomly assigned to sires. Hand crosses were performed daily, and all non-pollinated flowers were removed. After accounting for mortality in the greenhouse and planting logistics, the final pedigree consisted of 42 paternal half-sibling families and 124 maternal full-sibling families at Grey Cloud, 48 paternal half-sibling families and 132 maternal full-sibling families at McCarthy Lake, and 21 paternal half-sibling families and 60 maternal full-sibling families at CERA.

### Field planting design

In October-November of each of three consecutive years (2014-2016), pedigreed seeds from each respective population were planted into restored tallgrass prairie near to (<1 km) each site of origin. To maintain the environment for the experimental populations as realistic as possible, vegetation was left in place and only mown (or burned in the case of CERA in 2016 and McCarthy Lake in 2014) in the fall to facilitate planting. In Fall of 2014, all rows needed for all three years of the study were laid in parallel two meters apart, each row 50 m long. For each Fall planting, a subset of rows within blocks was chosen at random, and planting positions were marked with nails at 2 m intervals. Pedigreed seeds were planted at randomly chosen positions according to the following protocol. Seeds from each full-sibling family from the greenhouse crosses were combined and haphazardly distributed into envelopes, such that each envelope contained five seeds. We randomly assigned envelopes to positions along rows in a randomized block design (eight blocks at Grey Cloud and McCarthy Lake, and four blocks at CERA positioned in the same location in all years), resulting in 15-24 replications of each half-sib family per block. Blocks contained 25-26 rows (13 rows at McCarthy Lake). At each position centered at the nail, five seeds from a randomly chosen full-sib family were planted at 10 cm spacing, approximately 1 cm under the soil surface.

### Fitness surveys

We regularly recorded fitness components for each planted seed throughout the growing season of the following year, including emergence, survival to flowering, and reproductive output (Table 1). A plant was considered present if cotyledons had emerged, and survival to flowering was scored based on the presence of open flowers or evidence of flowering (e.g., wilted flowers, pedicels, or fruit). At each late-season census (i.e. after flowering), we recorded the number of ripe fruit collected, number of fully elongated but immature fruits collected, number of fruits collected off the ground in the immediate vicinity, number of immature fruits, and pedicels from dehisced fruits remaining on the plant (see Sheth et al. 2018). Herbivory, ranging from light browsing to consumption of the entire plant, or severing of the stem near the soil surface, was recorded throughout the growing season. When herbivory occurred early in the growing season (i.e. June-July), plants sometimes recovered and eventually reproduced. Therefore, we retained these plants in censuses after herbivory to account for this late season seed production. Because individuals ranged widely in number of fruits set (1-68), and fruits explosively dehisce at maturity, we were unable to collect all mature fruits or count all seeds each plant produced. Therefore, seed counts were obtained on a subsample of fruits produced. Subsampling was accounted for in the statistical analyses (see below).

**Table 1.** Range of dates for collection of fitness expression data, for three sites of *Chamaecrista fasciculata*, over three consecutive years.

|  | Year |  |  |
| --- | --- | --- | --- |
| Site | 2015 | 2016 | 2017 |
| Grey Cloud | June 1 – October 1 | June 10 – October 7 | June 5 – October 3 |
| McCarthy Lake | May 27 – September 25 | June 7 – October 11 | June 8 – October 5 |
| CERA | May 10 – September 23 | June 2 – September 29 | June 1 – October 23 |

### Statistical Analyses

Estimates of 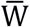 and V_A_(W) for each population in each year (Geyer et al. 2007; Shaw et al. 2008) we conducted aster analyses, as implemented in the package “aster” (Geyer 2018), using the R 3.5.1 environment (R Core Team 2018). The graphical model for lifetime fitness (Fig. 1) included seedling emergence and survival of plants to flowering, both of which were modeled as Bernoulli variables, and the number of fruits and seeds produced, which were modeled with Poisson distributions. To account for subsampling of fruits, we followed Appendix S1 of Stanton-Geddes et al. (2012a), incorporating in our graphical models an additional node representing the total number of fruits sampled. This leads to estimates that are scaled up to the total number of seeds produced per seed planted. Using this model for expression of components of fitness through the lifespan, as well as the dependence of each on components expressed earlier, we obtained unconditional estimates, i.e. the mean and additive genetic variance of the number of seeds produced per seed planted.

For each population in each year, we conducted separate aster analyses to estimate V_A_(W), including as predictors parental (genetic) effects and block. As in Sheth et al. (2018), only parental (genetic) effects were treated as random as the basis for estimating V_A_(W). Models including factors whose levels are considered random (i.e. for which the parameter of interest is variance of the levels’ effects) were introduced by Fisher (1918) with the motivation of estimating genetic variance of a quantitative trait. The original statistical methodology relies heavily on the convenient mathematical properties of the assumed Gaussian distribution of the random effects. Geyer et al. (2013) outline the theoretical and computational problems that attend relaxation of the Gaussian assumption for the random effects, especially with multiple random factors, in aster models and other GLMM, which are explicitly motivated by the need to model cases that do not meet that assumption (see also Bolker et al. 2009). We follow the recommendation of Geyer (2015, slide 3) to designate as random only those factors necessary to address the questions of scientific interest, here, the parental effects, enabling estimation of genetic variance for fitness. In addition, to test whether parental genetic effects differed among years, we jointly analyzed the data for cohorts grown in all three years, including year as a fixed factor and parental (genetic) effects and the interaction between parental effects and year (genotype-by-year) as random factors. We tested fixed factors with likelihood ratio tests of the full model vs. models excluding individual factors. To obtain estimates of additive genetic variance for lifetime fitness, we first conducted separate analyses with sire and dam as random factors, finding both significant and roughly comparable in magnitude. We then constructed models that explicitly equated the variance due to sires with that for dams (i.e. modeling the equal nuclear transmission of each parent to their offspring) and estimated a single variance component for their effects (hereafter parental effects), with sire effects being (see Sheth et al. 2018). From separate analyses, also including blocks as fixed effects, we estimated 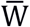.

For each population-year combination, mean fitness was represented by the estimate for mean fitness (with its standard error) that is the median value among the estimates of mean fitness for all the blocks of a given site. To visualize breeding values for fitness on the measurement scale (i.e. number of seeds produced per seed planted), we performed non-linear transformations of breeding values on the canonical parameter scale to the mean-value parameter scale using a mapping function that adds a component of a fixed effect (see Table 3 for relevant blocks) to each random parental effect (Geyer and Shaw 2013; de Villemereuil et al. 2016). The resulting values represent estimates of breeding values on the mean-value parameter (i.e. measurement scale). Estimates of V_A_(W) on the measurement scale were obtained via the delta method following Geyer (2019). We obtained asymmetrical 95% confidence intervals using the parametric bootstrap to obtain a bootstrap-t distribution from the difference between original and 100 bootstrapped V_A_(W) values divided by their standard error (DiCiccio and Efron 1996). Predicted changes in mean fitness from FTNS 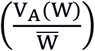 were added to the estimates of mean fitness 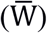 to generate the predicted mean fitness of the progeny generation. Standard errors were calculated for predicted changes in fitness 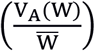 following Geyer (2019). Apart from the possibility of seed dormancy, our fitness records cover the entire life history of individual plants, and therefore represent estimates of total absolute fitness over individual life spans.

To compare the genetic effects on fitness (breeding values) between years, and the potential for temporal fitness trade-offs, we obtained breeding values for each sire in each year and plotted each sire’s value in one year against its breeding value in another for all three combinations of years. We also calculated Pearson’s product-moment correlation coefficients between inferred breeding values expressed in different years. We emphasize that these correlations do not account for the uncertainty associated with inferred breeding values; we present them only as a coarse description of association between breeding values expressed in different years. All data and scripts for the results reported here are available at https://github.com/mason-kulbaba/adaptive-capacity.git

## Results

### Expression of additive genetic variance for lifetime fitness

We found highly significant additive genetic variance for lifetime fitness, V_A_(W), in all three years for all three populations, as reflected by significant parental effects in all models (Table 2); block effects were also significant in all cases, indicating spatial variation within sites in expression of fitness. Estimates of V_A_(W) varied widely among years and populations, with the smallest and largest estimates of V_A_(W) found in consecutive years for the Grey Cloud population (V_A_(W) = 0.631 in 2016, and 6.491 in 2017). The statistical detection of V_A_(W) (Table 3) implies capacity for genetic response to natural selection based on standing V_A_(W). Probability distributions of breeding values for each of the nine cases are displayed in Fig. 2.

**Figure 2.**
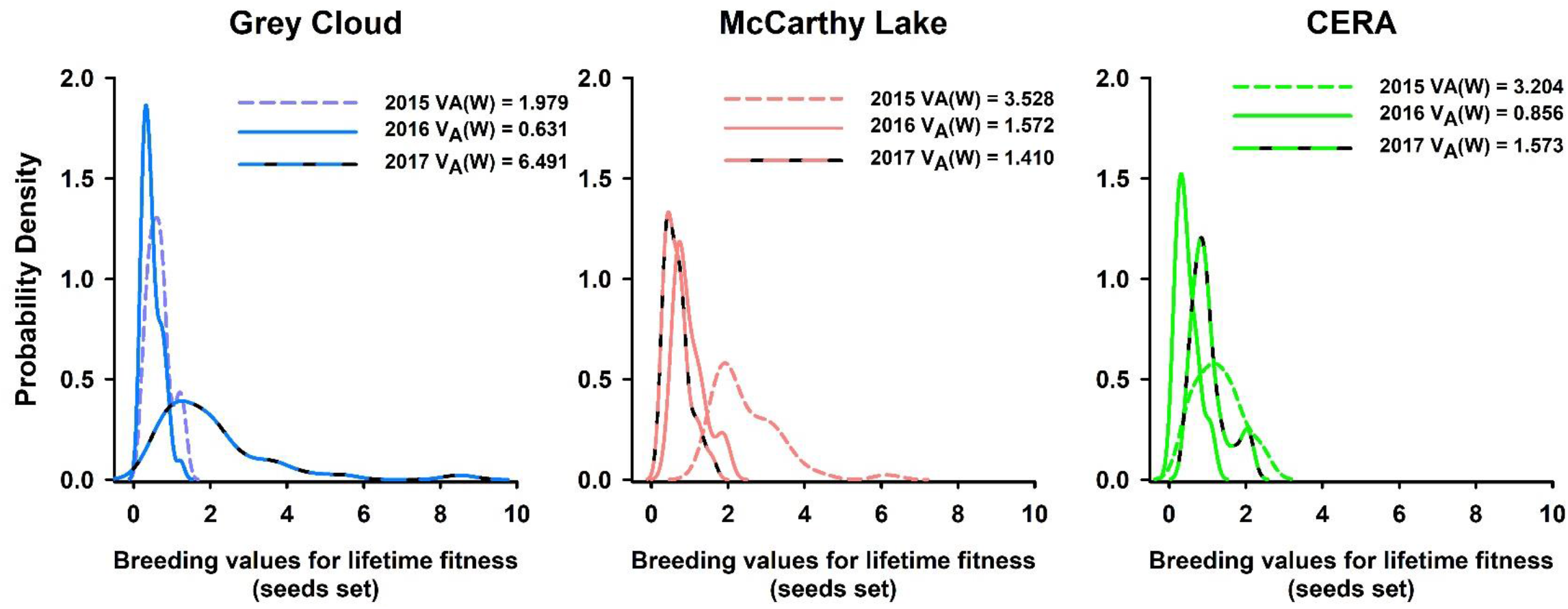
Additive genetic variation for lifetime fitness for three populations of *Chamaecrista fasciculata*, in three consecutive years. Lines represent probability density distributions of estimated breeding values for lifetime fitness (seeds set).

**Table 2.** Summary from aster models testing the fixed effects of block and year, the random factors of combined parental effects, and the interaction between parental effects and year on individual lifetime fitness. Statistical significance of predictor variables was assessed using likelihood ratio tests, and random parental effects were assessed from summary output of aster analyses.

| Model | Test df | Test deviance | P |
| --- | --- | --- | --- |
| Grey Cloud – *G x Year* model |  |  |  |
| Block | 23 | 159.91 | $P < 0.0001$ |
| Year | 2 | 47.28 | $P < 0.0001$ |
| Parental effects | -- | -- | $P < 0.0001$ |
| Year × Parental effects | -- | -- | $P < 0.0001$ |
| McCarthy Lake – *G x Year* model |  |  |  |
| Block | 23 | 159.91 | $P < 0.001$ |
| Year | 2 | 47.28 | $P < 0.001$ |
| Parental effects | -- | -- | $P < 0.0001$ |
| Year × Parental effects | -- | -- | $P < 0.0001$ |
| CERA – *G x Year* model |  |  |  |
| Block | 11 | 189.80 | $P < 0.0001$ |
| Year | 2 | 58.95 | $P < 0.0001$ |
| Parental effects | -- | -- | $P < 0.0001$ |
| Year × Parental effects | -- | -- | $P < 0.0001$ |

**Table 3.** Additive genetic variance for fitness (V_A_(W)) and 95% confidence intervals, mean fitness (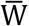 and standard error, predicted change in mean fitness 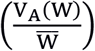 and standard error, and predicted mean fitness of second generation determined for each of three sites and three years of *Chamaecrista fasciculata*. Block used to estimate mean fitness indicated in parentheses after year.

| Site/Year<br>(Block) | Number of<br>Seeds<br>Planted | $V_A(W)$<br>(SE) | $\bar{W}$<br>(SE) | $\frac{V_A(W)}{\bar{W}}$<br>(SE) | Predicted Fitness<br>of Gen. 2 |
| --- | --- | --- | --- | --- | --- |
| Grey Cloud Dunes |  |  |  |  |  |
| 2015 (Block 6) | 3658 | 1.858<br>(0.558, 3.021) | 1.11<br>(0.17) | 1.67<br>(0.23) | 2.78 |
| 2016 (Block 6) | 3660 | 0.830<br>(0.193, 1.410) | 0.64<br>(0.13) | 1.30<br>(0.14) | 1.94 |
| 2017 (Block 7) | 3660 | 6.491<br>(1.365, 9.161) | 1.06<br>(0.34) | 6.11<br>(2.32) | 7.17 |
| McCarthy Lake |  |  |  |  |  |
| 2015 (Block 4) | 3445 | 3.528<br>(1.393, 16.503) | 3.02<br>(1.23) | 1.17<br>(0.89) | 4.19 |
| 2016 (Block 5) | 3480 | 1.572<br>(0.563, 5.103) | 1.88<br>(0.57) | 0.84<br>(0.18) | 2.72 |
| 2017 (Block 6) | 3516 | 1.410<br>(0.610, 2.157) | 1.08<br>(0.18) | 1.31<br>(0.18) | 2.39 |

| CERA |  |  |  |  |  |
| --- | --- | --- | --- | --- | --- |
| 2015 (Block 1) | 1845 | 3.204<br>(0.983, 6.922) | 1.80<br>(0.24) | 1.79<br>(0.65) | 3.58 |
| 2016 (Block 2) | 1750 | 0.856<br>(0.295, 2.301) | 0.73<br>(0.11) | 1.18<br>(0.13) | 1.91 |
| 2017 (Block 3) | 1750 | 1.573<br>(0.383, 3.694) | 1.24<br>(0.25) | 1.27<br>(0.49) | 2.51 |

Significant genotype-by-year interactions for each population (Table 2) indicated that the expression of parental genetic effects varied among years. Year-specific breeding values within each population revealed large differences between years in family-specific expressions of total lifetime fitness (Fig. 3). Breeding values exhibited small to moderate correlations between years, with two cases of modest negative correlations: 2015 vs. 2016 in McCarthy Lake and 2016 vs. 2017 in CERA (Fig. 3). The findings of significant genotype-by-year interactions indicate that families contributing disproportionately to the next generation often differ among years.

**Figure 3.**
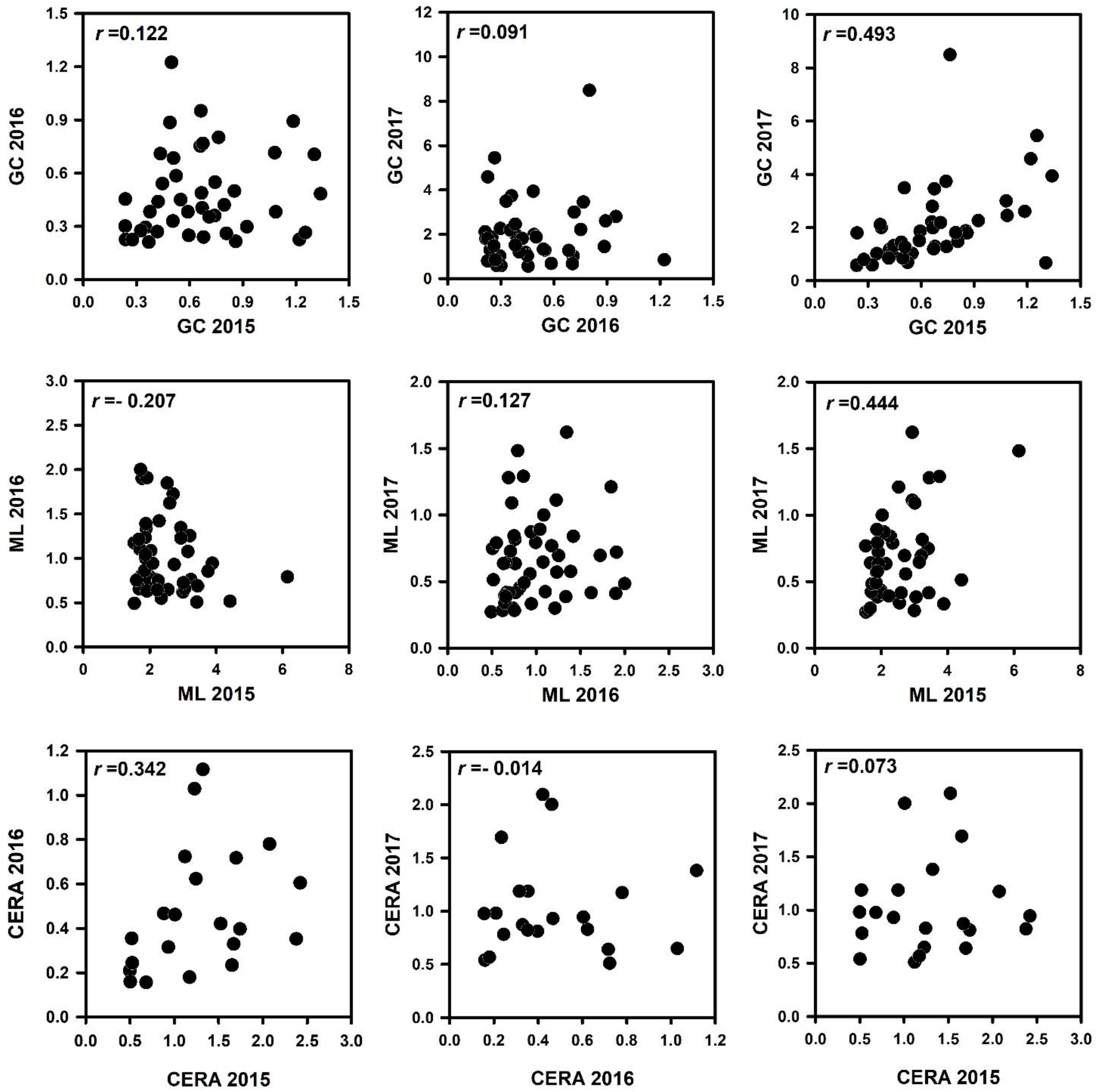
Paternal family-specific breeding values for lifetime fitness (seeds set) compared across three consecutive years in three populations of *Chamaecrista fasciculata.* Pearson correlation coefficients do not account for error associated with estimates of breeding values and are therefore presented as a coarse description of the interannual relationship between breeding values. Please note differences in scale across panels.

### Mean fitness and predicted changes in mean fitness

The inclusion of block effects in all fixed-effects aster models for 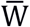 explained significantly more variation than models without (Grey Cloud and McCarthy Lake: all test df = 7, all test deviance > 4.26, all P < 0.05; CERA: all test df = 3, all test deviance > 27.92, all P < 0.0001).

As with estimates of V_A_(W), estimates of mean fitness varied among populations and years (Table 3; Fig. 4). Mean fitness was highest in the McCarthy Lake site in 2015 (mean and standard error: 3.528 ± 1.227) and lowest in Grey Cloud in 2016 (0.640 ± 0.129). This and one other instance (CERA mean and standard error: 0.725 ± 0.112) indicating declining populations (i.e. 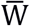) were found in the same year.

**Figure 4.**
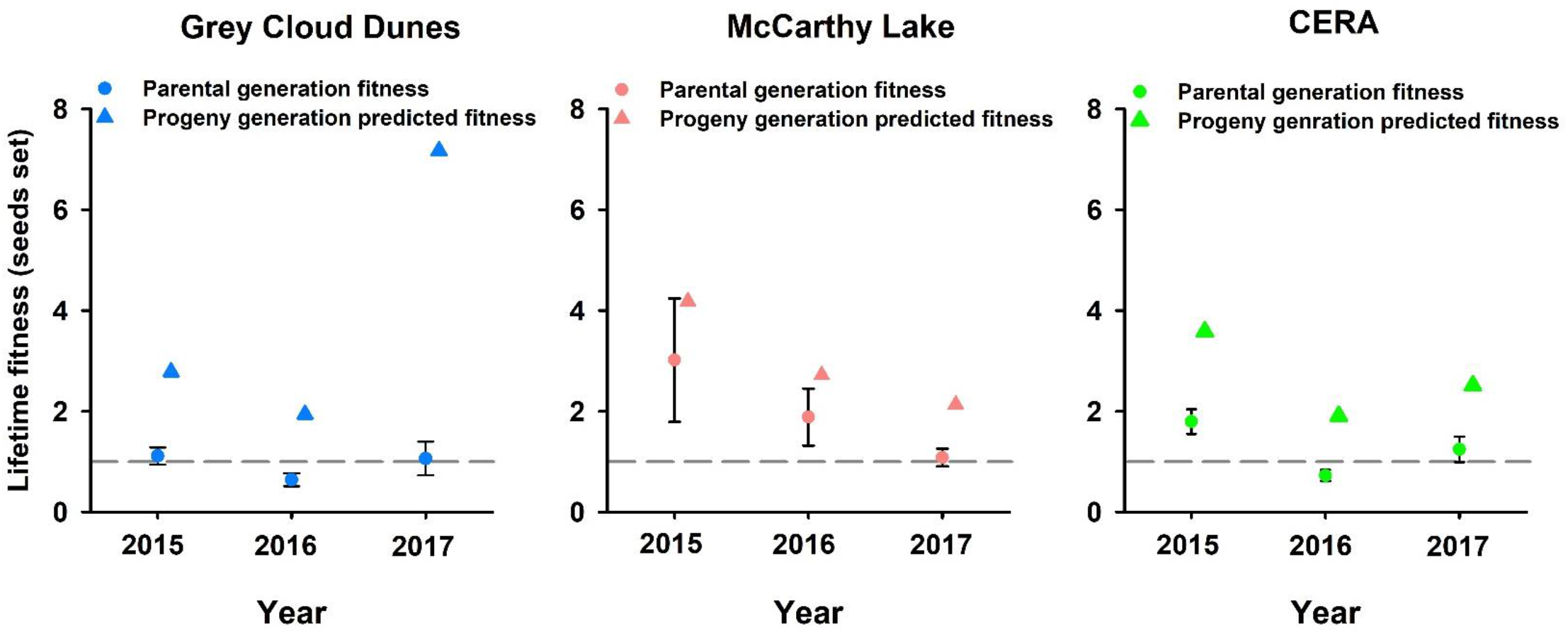
Estimated mean fitness of parental (circles with standard errors) and progeny generations (triangles with standard errors) for three populations of *Chamaecrista fasciculata* in three consecutive years. Horizontal dashed lines represent mean fitness of 1, indicating individual replacement and population stability.

The estimates predicting change in mean fitness from Fisher’s Fundamental Theorem 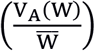, are biologically meaningful, ranging from increases of just under 1 seed per individual seed planted to about 6, varying with the magnitude of both 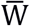 and V_A_(W). The largest increase in mean fitness was predicted for the progeny of Grey Cloud plants after 2017 (mean increase of 6.122 seeds per plant) and the smallest for progeny of McCarthy Lake plants after 2016 (mean increase of 0.834 seeds per plant). Notably, we identified two predictions of evolutionary rescue for the progeny generations of 2016 in Grey Cloud and CERA. This predicted increase in mean fitness is consistent with population sustainability and growth of the progeny generations (Table 3; Fig. 4).

## Discussion

The immediate capacity for population-level adaptation is strongly dependent on the presence and magnitude of additive genetic variance for lifetime fitness (Fisher 1930; Ewens 2004). Our study detected significant and substantial capacity for immediate adaptation through current natural selection on standing levels of V_A_(W) in three populations in three years. Among these nine cases, we detected two instances of populations declining numerically, and in those cases obtained predictions of evolutionary rescue increasing mean fitness to levels consistent with maintaining or increasing population size. Genotype-by-year interactions indicated that genetic effects on fitness differed among years. These interactions reflect differences among years both in the magnitude of V_A_(W) and in genotypic fitness rankings. Predicted change in mean fitness also varied among years, suggesting that the capacity for adaptation is strongly influenced by temporal environmental variation. Below, we discuss the importance of these findings and consider how our results from direct study of the adaptive process in the wild illuminate the potential for ongoing adaptation. We also relate our findings to the potential for evolutionary rescue of declining populations. Finally, we describe how our experimental and analytical approaches overcome obstacles commonly associated with evaluating adaptive capacity in the wild, and we advocate for the broader implementation of these approaches.

### Expression of additive genetic variance for lifetime fitness

Our detection of prevalent and non-negligible additive genetic variance for lifetime fitness, characterized as the number of seeds produced per seed planted, indicates differential genetic contributions to the progeny generation and, via the FTNS, clear capacity for ongoing adaptation under natural selection in each year. We estimated significant V_A_(W) in the home site of all three populations in all three years of study (Fig. 2), albeit at varying magnitudes. These results contrast with those of several studies that conclude limited capacity for adaptation, through estimates of heritability of fecundity (e.g. Kruuk et al. 2000), or other individual fitness components (Mousseau and Roff 1987; Matos et al. 2000; McCleery et al. 2004; Teplitsky et al. 2009). We suggest that these studies can be reconciled with our findings in part by recognizing that even when additive genetic variances of individual fitness components are modest, estimates of V_A_(W) may be considerably larger due to the compounding of fitness components through an individual’s life-history (Shaw et al. 2008; Shaw and Geyer 2010). This compounding plays out over the entire life-history, regardless of the extent to which there are genetically based tradeoffs between individual components of fitness (e.g. Rose and Charlesworth 1981). Our analysis of total lifetime fitness through aster modeling enabled us to account for the dependent nature of sequential fitness components in our estimates of total fitness. V_A_(W) may also be obscured by environmental variation (Price and Schluter 1991).

Our finding of prevalent interactions between genotype and year can be partitioned into two aspects of environmental dependence of genetic expression (Falconer 1952). First, correlations between years of family-specific breeding values for lifetime fitness were generally low and, in two cases, slightly negative (Fig. 3). Thus, genotypic contributions of offspring in one year are not predictive of contributions in another year. Interestingly, the two strongest genetic correlations between years were not for consecutive years. These modest between-year genetic correlations suggest that the response to selection may not be accompanied by directional change in the frequency of the same alleles over multiple years. Second, for each population, estimates of V_A_(W) differed strikingly among years (Table 3), though we acknowledge substantial uncertainty in the estimates. Nevertheless, these differences in estimated V_A_(W) suggest that the immediate capacity for ongoing adaptation varies among years. Almost certainly, many aspects of environment, whether slight or major, differed among the years at each site; to determine which of these influenced genetic disparities in fitness would require experimental manipulation of particular aspects of the environment, as conducted by Torres-Martinez et al. (2019).

Whereas we found considerable V_A_(W) for multiple populations, explanations for the maintenance of such potential variation has long been elusive (e.g. Burt 1995). One potential explanation is that trade-offs between individual fitness components maintain additive genetic variation (Barton and Keightley 2002). Our previous work focusing on the Grey Cloud population revealed positive correlations of breeding values between fitness components and also two environments (Sheth et al. 2018). This result suggests that trade-offs (i.e. negative genetic correlations) do not account for the maintenance of observed levels of V_A_(W).

Temporal environmental variation has long been conjectured to maintain genetic variation, but the theoretical conditions that predict this outcome are restrictive (Felsenstein 1976). Experiments manipulating environmental variation have led to conflicting conclusions on the role of environmental variation in the maintenance of genetic variation (Mackay 1980; Yeaman et al. 2010; Huang et al. 2015). Through estimates of total lifetime fitness, instead of individual traits or components of fitness, our results have documented extensive temporal variation in the expression of breeding values within populations (Fig. 2; Fig. 3). Given the short timeframe of our study (i.e. single generation replicated over three years), the impact of novel mutations on lifetime fitness would be negligible. Additional phenomena may also contribute to the conservation of genetic variation (e.g., marginal overdominance; Levene 1953; Gillespie 1984), but the differential and variable genetic contribution of families to the following progeny generation across years (Fig. 3) is consistent with a role for temporal environmental variation in impeding depletion of V_A_(W) by natural selection. These annual variations may contribute to the maintenance of the appreciable levels of observed V_A_(W) in our study.

Our measures of immediate capacity for adaptation can alternatively be expressed as evolvability *(sensu* Houle 1992, see Table 3) to yield the change in absolute mean fitness as a proportion of the current absolute mean fitness. This measure of predicted proportional change has been advocated particularly in the context of trait evolution. In the context of predicting change in mean fitness, we view the original formulation of the FTNS as more informative because it is directly interpretable demographically as the predicted change in the *per capita* contribution of offspring to the population. In contrast, there is no direct demographic interpretation of the corresponding evolvabilities.

### Mean fitness and predicted changes in mean fitness

Similar to estimates of V_A_(W), mean fitness and the predicted rate of adaptation across generations varied with population and year. Predictions based on Fisher’s Fundamental Theorem 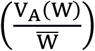 indicated that even the more modest estimates of V_A_(W) (e.g., Grey Cloud and CERA in 2016; Table 3) were sufficient to increase mean fitness under contemporary natural selection. In the case of the largest estimate of V_A_(W) (Grey Cloud in 2017), contemporary selection was predicted to drastically increase fitness to several times that of the parental generation (Table 3, Fig. 4).

Our study identified two cases where evolutionary rescue was predicted to increase mean fitness of declining populations to levels consistent with sustainability and even population growth. Mean fitness of both the Grey Cloud and CERA populations during the 2016 growing season was less than 1.0, indicating declining numbers of individuals to the following year. The prediction of genetic change in mean fitness, however, is that fitness in the progeny generation would increase to more than double the current mean fitness. Realization of this prediction would not only prevent further population decline but would increase seed production to result in population growth. Whereas evolutionary rescue has been directly demonstrated in controlled laboratory systems (Bell and Gonzalez 2009, 2011), examples from wild systems are lacking. Beginning with the work of Gomulkiewicz and Holt (1995), mathematical modeling efforts have described the likelihood of evolutionary rescue under various demographic and environmental scenarios, but the promise of evolutionary rescue of declining and threatened populations remains unclear (Bell 2017), as does their long-term sustainability. Therefore, along with Gomulkiewicz and Shaw (2012) we advocate for further empirically-based predictions of adaptation, as presented in our study, to elucidate the potential for evolutionary rescue in the wild.

Our assessments of fitness encompass components of fitness expressed across the entire life span of individuals, from each seed planted through to the seeds it produced; these fitness evaluations are thus uncommonly complete. Nevertheless, some aspects of fitness are not included. For example, this study did not account for fitness realized through siring of seeds (male fecundity). In a companion study, however, Kulbaba and Shaw (in review) found positive genetic correlations between lifetime fitness measured as maternal contributions (seeds set) and lifetime fitness measured through paternal contributions (seeds sired), regardless of population density and genetic relatedness among individuals. We therefore expect paternal-specific fitness to scale proportionately with maternal-specific fitness. A second omission from our fitness estimates is the potential for a delayed contribution to fitness due to seed dormancy. Partial surveys of germination in later years found low emergence from seeds planted earlier than the previous Fall, representing a modest proportion of each cohort. Moreover, in increasing populations, (i.e. 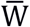 > 1), as we found in most cases, delays in an individual’s life-cycle are expected to have a discounting effect on its contributions to fitness (Roughgarden 1996, Eqs. 18.31, p. 331). On the temporal scale of adaptation of annual populations between consecutive years, which is our present focus, it is valid to neglect such delays. To address ongoing adaptation on longer time scales, further studies that include instances of delayed germination in estimates of fitness, with appropriate discounting, as in Eck et al. (2015) would be worthwhile. This is especially so, in view of our finding of weak correspondence of genetic selection between consecutive years (Fig. 3).

Our study has generated precise quantitative predictions of genetic change in mean fitness. However, correspondence between predicted genetic change in mean fitness across generations and the change in mean fitness that is realized remains to be assessed. We are currently evaluating this correspondence to address the accuracy of predictions of the rate of adaptation from Fisher’s FTNS for these populations.

### Conclusions

Whereas numerous authors invoke relentless selection to explain limited heritability of total fitness, or traits closely related to fitness (Kruuk et al. 2000; Coltman et al. 2005; McFarlane et al. 2014; de Villemereuil et al. 2019), our results demonstrate the persistence of high V_A_(W) despite contemporary selection. Significant interactions between genotype and year reflected modest correlations between years of family-specific breeding values for lifetime fitness, as well as estimates of V_A_(W) that varied widely across years. Along with estimates of V_A_(W), mean absolute fitness varied among sites and years. Using Fisher’s Fundamental Theorem, we predicted the rate of adaptation, and two instances of evolutionary rescue from standing V_A_(W) under contemporary natural selection. Our results reveal the power and utility of Fisher’s Fundamental Theorem to identify the capacity for adaptation, and to generate quantitative predictions for fitness increase in the wild.

Our experimental approach in combination with aster analyses for total lifetime fitness provided a direct estimate of V_A_(W) and fitness change (Shaw 2019). Many studies concerning the potential for adaptation rely on indirect estimates of fitness, and often focus on one or a few individual traits or fitness components, thus omitting the role of multiple life-history fitness expressions. Our approach to evaluating total lifetime fitness with aster models provides a more complete picture of fitness and V_A_(W). We recommend our empirical and analytical strategy for future studies of other organisms having different life-histories to more fully characterize the capacity for ongoing adaptation.

## Notes

#### Summary of Updates

minor change in analyses that produce asymmetrical 95% confidence intervals for estimates of VA(W), and standard errors for the predicted change in mean fitness. Also includes numerous wording changes to address questions of evolvabilities and evolution of traits.

https://github.com/mason-kulbaba/adaptive-capacity.git

